# Vascular mimicry by VE-cadherin enables trophoblast endovascular invasion and spiral artery remodeling during placental development

**DOI:** 10.1101/2022.02.15.480503

**Authors:** Derek C. Sung, Xiaowen Chen, Mei Chen, Jisheng Yang, Susan Schultz, Apoorva Babu, Yitian Xu, Siqi Gao, TC Stevenson Keller, Patricia Mericko, Michelle Lee, Ying Yang, Joshua P. Scallan, Mark L. Kahn

## Abstract

During formation of the mammalian placenta trophoblasts invade the maternal decidua and remodel spiral arteries to bring maternal blood into the placenta. This process, known as endovascular invasion, is thought to involve the adoption of functional characteristics of vascular endothelial cells (ECs) by trophoblasts through a process termed vascular mimicry. The genetic and molecular basis of vascular mimicry remains poorly defined, however, and whether trophoblasts utilize specialized endothelial proteins in an analogous manner to create vascular channels remains untested. Vascular endothelial (VE-)cadherin is a homotypic adhesion protein that is expressed selectively by ECs in which it enables formation of tight vessels and regulation of EC junctions. VE-cadherin is also expressed in invasive trophoblasts and is a prime candidate for a molecular mechanism of vascular mimicry by those cells. Here, we show that the VE-cadherin is required for trophoblast migration and endovascular invasion into the maternal decidua in the mouse. VE-cadherin deficiency results in loss of spiral artery remodeling that leads to decreased flow of maternal blood into the placenta, fetal growth restriction, and death. These studies identify a non-endothelial role for VE-cadherin in trophoblasts during placental development and suggest that endothelial proteins may play functionally unique roles in trophoblasts that do not simply mimic those in ECs.

## Introduction

During placental development in mice and humans, fetal trophoblasts invade the maternal decidua by a process known as endovascular invasion to remodel and connect to maternal spiral arteries^1–3^. This connection allows the flow of maternal blood through trophoblast-lined sinuses characteristic of hemochorial placentation^4,5^. Shallow trophoblast invasion, deficient spiral artery (SA) remodeling, and poor remodeling of the maternal decidua are features of placental dysfunction such as preeclampsia, a hypertensive condition of pregnancy that can lead to maternal and fetal complications^6,7^. Despite the broad and clinically significant impacts of placental dysfunction, the mechanisms controlling trophoblast endovascular invasion and SA remodeling remain poorly defined in vivo.

Invasive trophoblasts are believed to adopt an endothelial-like state by expressing endothelial specific genes^8–10^, a process that has been termed ‘vascular mimicry’ (reviewed in ^5^). Invasive trophoblasts in human and mouse placentas express vascular endothelial (VE)-cadherin (gene name Cdh5) during remodeling of SAs^9–11^. Invasive trophoblasts in preeclamptic placentas lack VE-cadherin^11^, and loss of VE-cadherin reduces trophoblast invasion in vitro^12^. These studies suggest a functional role for VE-cadherin in endovascular invasion and vessel formation. VE-cadherin is a well-studied cell-cell adhesion protein in the vascular endothelium where it regulates vascular integrity and growth and endothelial barrier function^13–15^. In vitro studies have suggested that VE-cadherin may regulate trophoblast-endothelial interactions^16^, but the requirement for VE-cadherin in trophoblasts during placental development in vivo remains unknown.

In the present study, we functionally tested the role of VE-cadherin in trophoblasts during placental development in mice. We find that conditional deletion of VE-cadherin from trophoblasts disrupts trophoblast invasion into the decidua and SA remodeling, resulting in placental insufficiency and fetal growth restriction. We show that VE-cadherin is important for trophoblasts to interact with and displace SA endothelium. These studies identify a molecular mechanism by which fetal trophoblasts use VE-cadherin to invade and remodel the maternal environment for successful pregnancy that is relevant to preeclampsia pathogenesis. They also provide a first functional test of the concept of trophoblast vascular mimicry in vivo, and suggest that canonical vascular endothelial proteins may be used by vascular trophoblasts in the placenta in ways that are specific for their function and do not merely mimic endothelial cell use.

## Results

### Trophoblast-specific deletion of VE-cadherin restricts placental and fetal growth and causes embryonic lethality

To understand the role of VE-cadherin in trophoblasts we generated Cyp19^Cre^; Cdh5^fl/fl^ (‘Cdh5 knockout’) placentas and mice in which VE-cadherin (encoded by Cdh5) is deleted specifically in fetal trophoblasts. Immunostaining for VE-cadherin and the trophoblast marker CK8 demonstrated efficient deletion of VE-cadherin in trophoblasts but not ECs in the Cdh5 knockout placenta (**Figure 1-Figure Supplement 1**). Cdh5 knockout embryos were present at the expected Mendelian ratio at E10.5-12.5 (**Figure 1-Table Supplement 1**, P=0.7667). However, there was an almost complete loss of trophoblast Cdh5 knockout embryos at E14.5-E16.5 and postnatal day 21 (**Figure 1-Table Supplement 1**, P<0.05 and P<0.005, respectively). Examination of placentas at E12.5 revealed that trophoblast Cdh5 knockout placentas were smaller and paler than those of control littermates in the same uterus (**Figure 1A-C**). Cdh5 knockout embryos exhibited marked fetal growth restriction and hemorrhaging at E12.5 (green arrowheads, **Figure 1D-F**). Histological and immunofluorescence analysis of E12.5 knockout embryos showed growth defects in numerous organs, including the heart, brain, and liver (**Figure 1-Figure Supplement 2**). These data demonstrate that trophoblast-specific loss of VE-cadherin confers fetal growth restriction and lethality. Importantly, since Cyp19^Cre^ activity is present only in placental trophoblasts^17,18^, these embryonic defects are secondary to placental defects.

**Figure 1.**
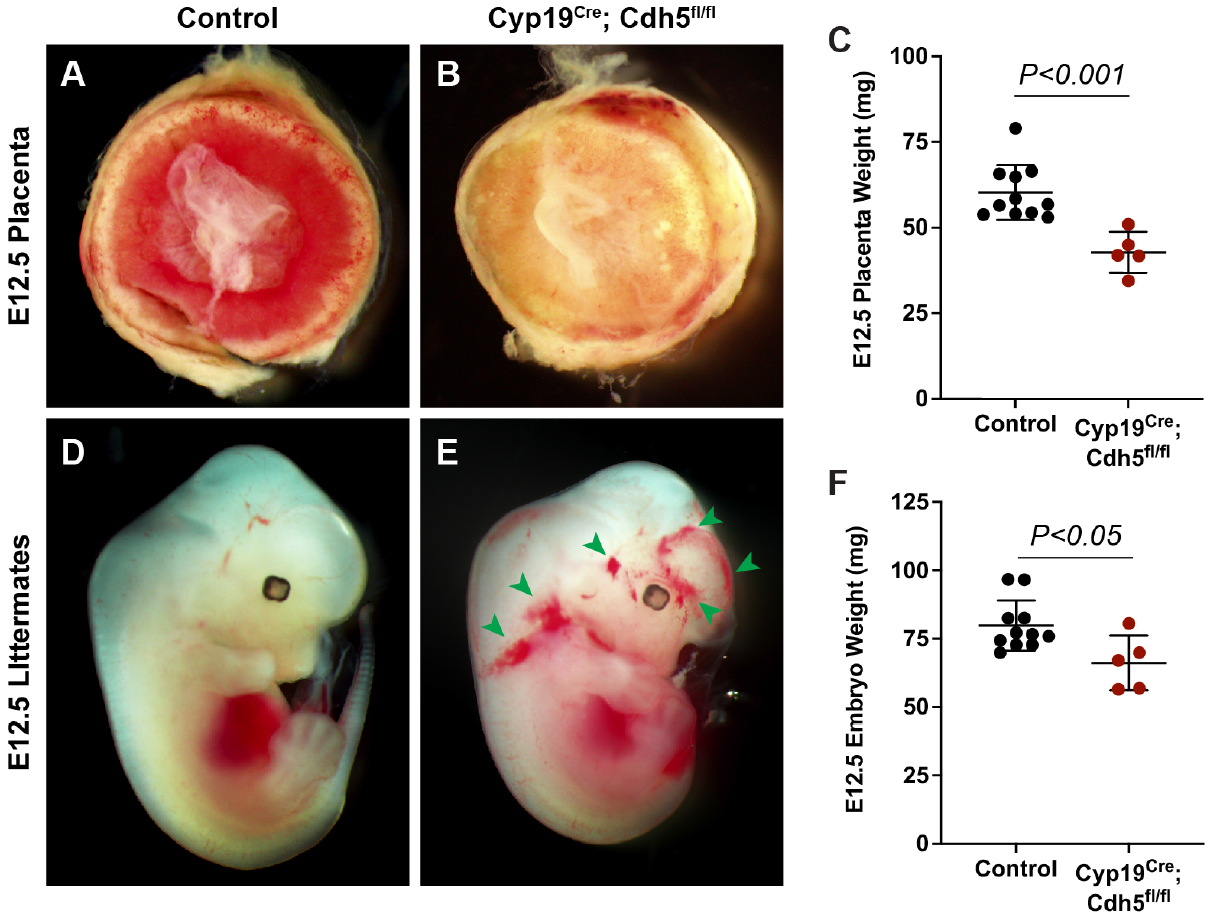
Deletion of VE-cadherin in trophoblasts results in placental and fetal growth restriction. **A-C,** Gross examination and quantification of E12.5 Control and Cyp19^Cre^; Cdh5^fl/fl^ placentas and weights. Control n=11, Cyp19^Cre^; Cdh5^fl/fl^ n=5. **D-F,** Gross examination and quantification of E12.5 Control and Cyp19^Cre^; Cdh5^fl/fl^ embryos and weights. Control n=11, Cyp19^Cre^; Cdh5^fl/fl^ n=5. Statistical analysis was performed using two-tailed, unpaired Welch’s t-test. Data are shown as means±S.D.

### Loss of trophoblast VE-cadherin disrupts endovascular invasion but not formation of the fetal vasculature

To characterize the placental defects conferred by trophoblast loss of VE-cadherin we performed histological staining with H&E and immunofluorescence staining for CK8 (trophoblasts), Endomucin (endothelial cells), and TER119 (red blood cells) on serial control and Cdh5 knockout placenta sections (**Figure 2A-D**). H&E and immunofluorescence staining both showed abundant trophoblasts, marked by CK8 positivity, surrounding Endomucin^+^ SAs within the decidua in control placentas (red arrowheads, **Figure 2B, B’**). Notably, trophoblasts surrounding maternal SAs were absent in knockout placentas (**Figure 2D, D’**), and quantification of number of trophoblasts and invasion depth into the decidua showed fewer and shallower invasion of trophoblasts overall (decidua, **Figure 2E, F**).

**Figure 2.**
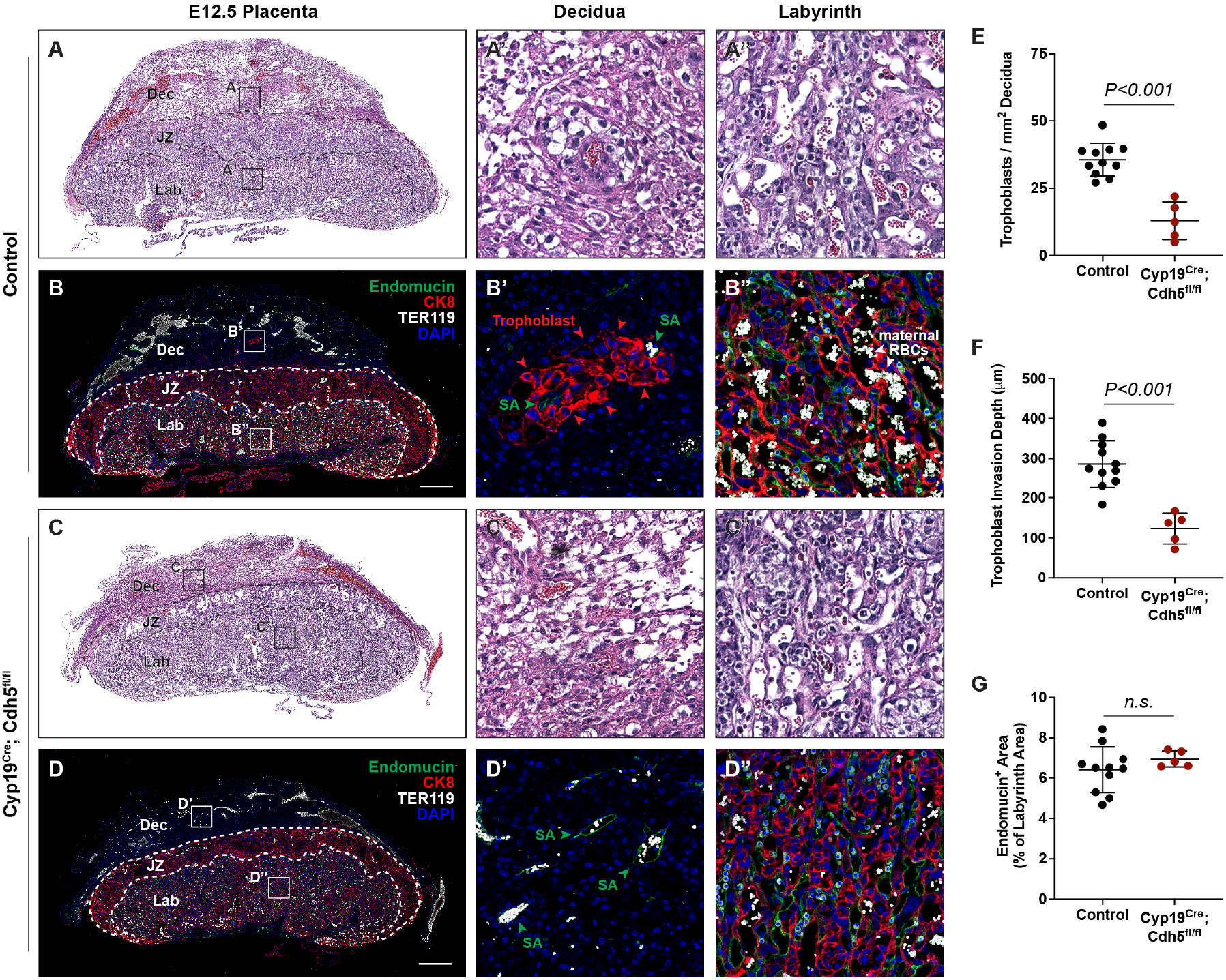
VE-cadherin promotes trophoblast endovascular invasion. **A-D,** Hematoxylin and eosin (H&E) staining and immunofluorescence staining for Endomucin (green), CK8 (red), and TER119 (gray) of E12.5 Control (**A, B**) and Cyp19^Cre^; Cdh5^fl/fl^ (**C, D**) serial placenta sections. Dotted lines demarcate the different placental regions. Red arrowheads indicate trophoblasts. Green arrowheads indicate spiral arteries. White arrowheads indicate maternal red blood cells (RBCs). Note fewer non-nucleated, maternal TER119^+^ cells in the labyrinth region of Cyp19^Cre^; Cdh5^fl/fl^ placentas. Boxes on the left correlate with magnified images on the right, and boxes in H&E and immunofluorescence images are of the same region. Scale bars = 500μm. Dec (decidua), JZ (junctional zone), Lab (labyrinth). **E-G,** Quantification of number of trophoblasts in the decidua (**E**), trophoblast invasion depth (**F**), and percent labyrinth Endomucin^+^ area (**G**). Control n=11, Cyp19^Cre^; Cdh5^fl/fl^ n=5. Statistical analysis was performed using two-tailed, unpaired Welch’s t-test. Data are shown as means±S.D.

The findings described above suggested that Cdh5 knockout placentas were less able to carry maternal blood to nourish the growing embryo. Fetal and maternal RBCs can be differentiated by the presence of nuclei in fetal RBCs. The labyrinth in knockout placentas had fewer enucleated TER119^+^ RBCs, indicating less maternal blood and consistent with paler placentas observed by gross examination (white arrowheads, **Figure 2B” vs. D”**). To determine whether loss of VE-cadherin affects the fetal placental vasculature, we quantified Endomucin^+^ vascular area in the labyrinth. We detected no differences in Endomucin^+^ staining, demonstrating that loss of VE-cadherin from trophoblasts does not disrupt formation of the fetal placental capillary plexus (**Figure 2B, D, G**). These findings suggest that VE-cadherin is important for trophoblast migration and for the association with SAs required to channel maternal blood into the placenta.

### Loss of trophoblast VE-cadherin blocks displacement of spiral artery endothelial cells and spiral artery remodeling

A critical early step in establishing maternal circulation to the placenta is trophoblast invasion of the decidua and its SAs. The observation that there were fewer VE-cadherin-deficient trophoblasts adjacent to SAs and decreased maternal blood within the labyrinth suggested that trophoblast VE-cadherin may play a requisite role in SA remodeling. Since loss of vascular smooth muscle is a key step in SA remodeling, we stained control and knockout placentas for alpha-smooth muscle actin (aSMA). SAs in Cdh5 knockout placentas exhibited persistent vascular smooth muscle coverage compared with those in control placentas (white arrowheads, **Figure 3A, B**). These studies reveal that loss of trophoblast VE-cadherin disrupts SA remodeling, which likely contributes to reduced maternal blood within the placenta.

**Figure 3.**
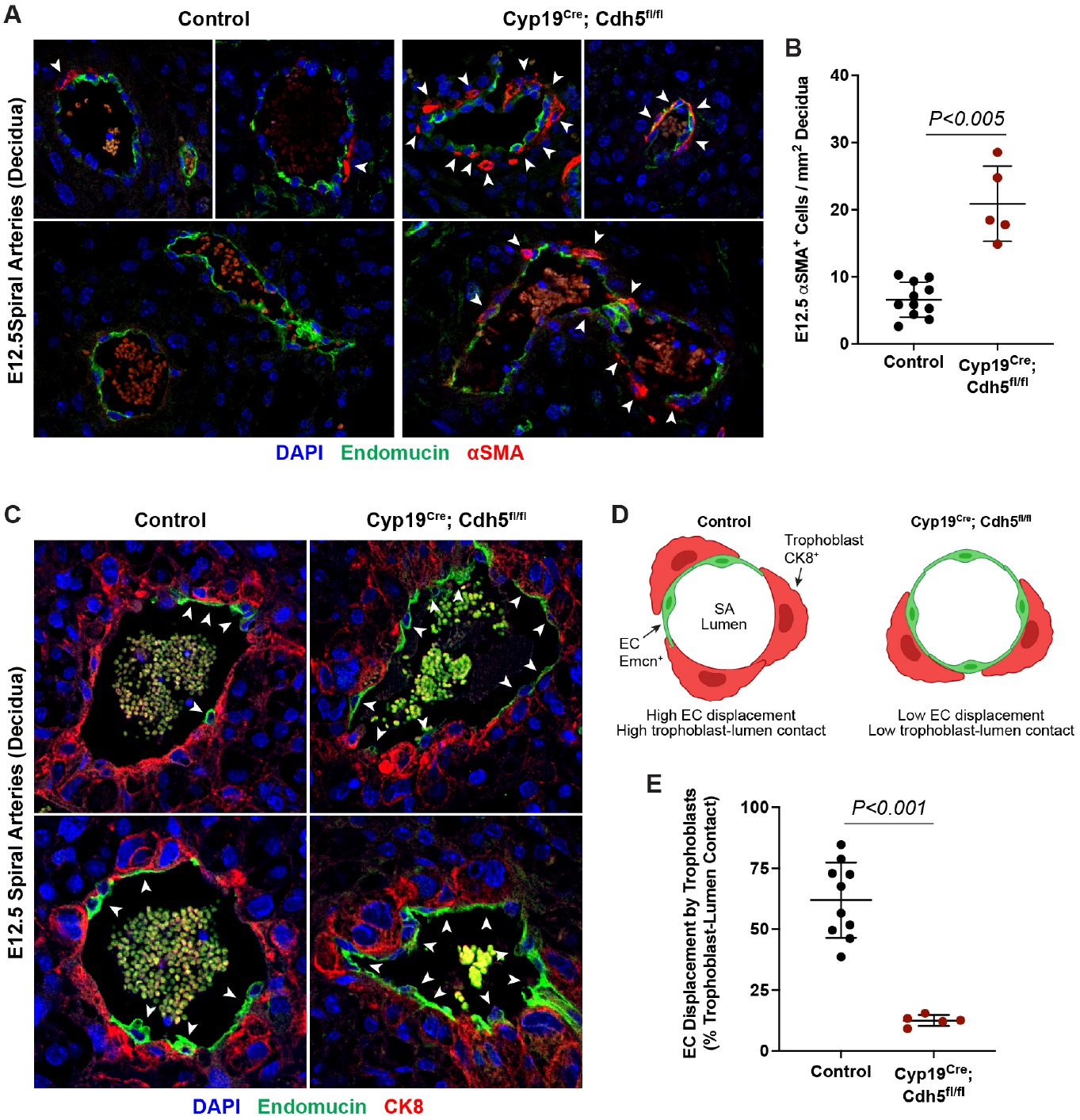
VE-cadherin is required in trophoblasts to remodel spiral arteries and the displace spiral artery endothelium. **A,** Immunofluorescence staining of E12.5 Control and Cyp19^Cre^; Cdh5^fl/fl^ placentas for Endomucin (green) and aSMA (red). White arrowheads indicate aSMA^+^ cells. **B,** Quantification of aSMA^+^ cells in decidua. Control n=11, Cyp19^Cre^; Cdh5^fl/fl^ n=5. **C,** Immunofluorescence staining of E12.5 Control and Cyp19^Cre^; Cdh5^fl/fl^ placentas for Endomucin (green) and CK8 (red). White arrowheads indicate Endomucin^+^ spiral artery endothelial cells. **D,** Schematic demonstrating differences in trophoblast and spiral artery (SA) endothelial cell (EC) contact with the vessel lumen. **E,** Quantification of the percent trophoblast-lumen contact, which was calculated by measuring the circumference of the vessel lumen and then measuring the length of CK8^+^ trophoblasts in contact with the lumen. Each point represents the average of at least 3 spiral arteries from an individual placenta. Control n=10, Cyp19^Cre^; Cdh5^fl/fl^ n=5. Statistical analysis was performed using two-tailed, unpaired Welch’s t-test. Data are shown as means±S.D.

During the process of SA remodeling, invasive trophoblasts displace the endothelial layer of SAs to direct maternal blood flow through trophoblast-lined sinuses into the labyrinth. In vitro studies have suggested that trophoblast expression of VE-cadherin may enable these cells to adhere to SA endothelial cells (ECs)^16^. The finding that trophoblast Cdh5 knockout placentas fail to remodel SAs suggested that there may be defects in trophoblast-SA interactions. To address the role of VE-cadherin at the site of trophoblast-SA connection, we first sought to image the site at which trophoblasts connect to SAs. Close inspection of this region in control placentas revealed a clear demarcation from luminal Endomucin^+^ SA endothelium to luminal CK8^+^ trophoblasts (white arrowheads, **Figure 3C, D**). In contrast, we found that SAs in VE-cadherin knockout placentas maintained a layer of intact ECs despite being surrounded by trophoblasts (white arrowheads, **Figure 3C, D**). Quantification of the percent of trophoblasts immediately in contact with the lumen demonstrated that knockout placentas had a lower percentage of trophoblasts and higher percentage of ECs covering the lumen compared to controls (**Figure 3E**). Together, these data suggest that VE-cadherin is required for trophoblast displacement of spiral artery ECs during endovascular invasion for efficient maternal-fetal circulatory connection.

### Defective trophoblast invasion and spiral artery remodeling cause placental insufficiency and fetal distress

The finding that Cdh5 knockout placentas have less maternal blood and are associated with mid-gestation embryonic lethality suggested that failed SA remodeling restricts maternal blood flow into the placenta, thus causing fetal demise. Human placental insufficiency is typically assessed with ultrasound measurement of placental hemodynamics and fetal heart rate, a readout for overall fetal health. We therefore utilized Doppler ultrasound to measure peak systolic and end diastolic velocities in the umbilical arteries (**Figure 4A**) and calculated resistance and pulsatility indices (RI and PI) and fetal heart rates to assess placental vascular resistance and fetal wellbeing in individual concepti. Elevated RI and PI values are clinical indicators of placental insufficiency in humans and are associated with conditions such as preeclampsia and fetal growth restriction. Fetal heart rate is used as a clinical parameter for fetal wellbeing, with fetal bradycardia indicative of fetal distress. While control embryos had RIs, PIs, and fetal heart rates within range of previously published studies^19^, we found that trophoblast Cdh5 knockout embryos exhibited significantly increased RI and PI (**Figure 4B-D**) and significantly reduced fetal heart rates (**Figure 4B, E**), consistent with placental insufficiency and fetal distress. Knockout embryos also exhibited reversal of end-diastolic flow as shown by the directional change of velocity from negative (peak systole) to positive (end of diastole) (**Figure 4B**), indicative of high vascular resistance. These hemodynamic data demonstrate placental insufficiency that contributes to fetal growth restriction and fetal demise following loss of trophoblast invasion and spiral artery remodeling.

**Figure 4.**
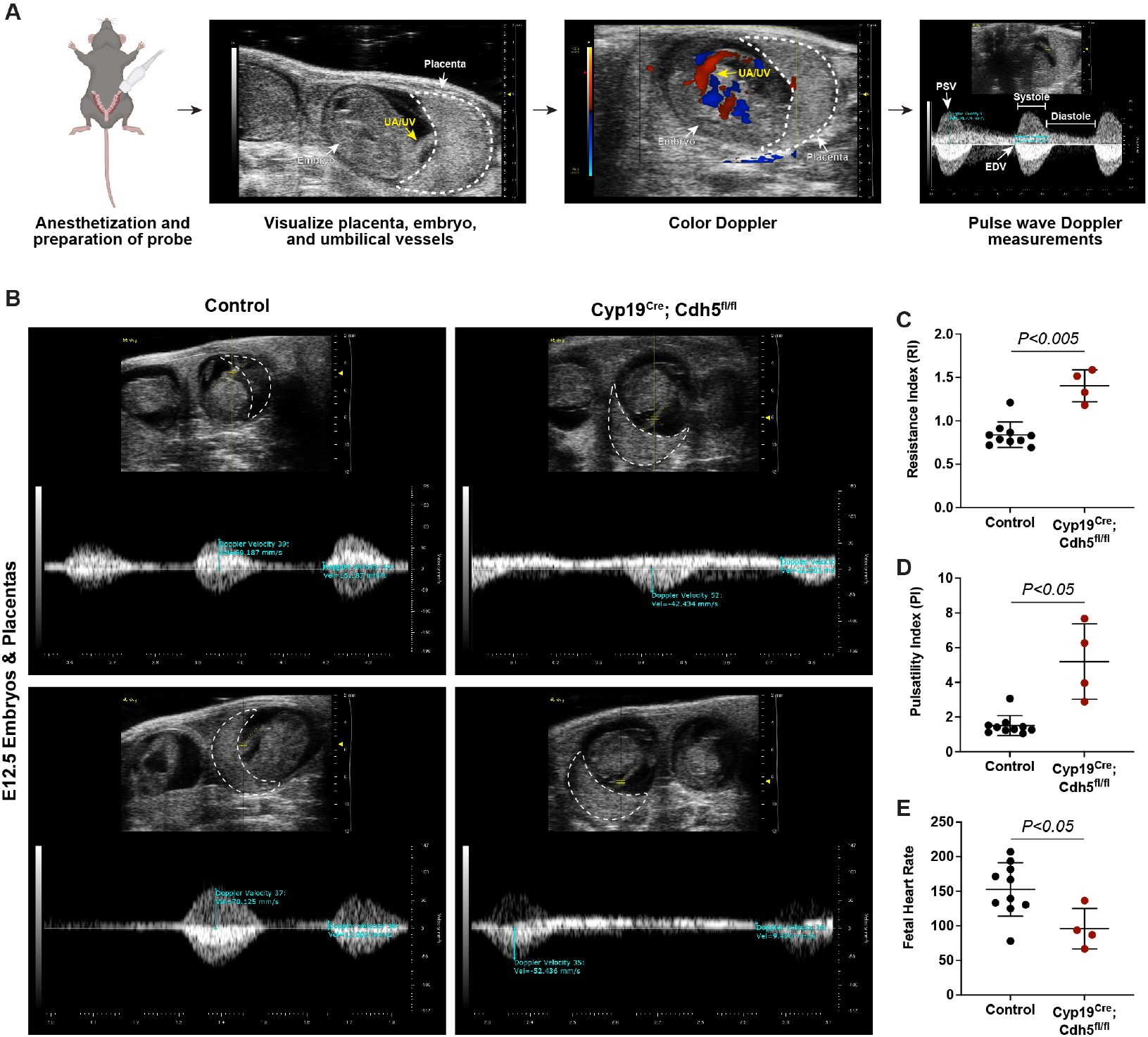
Defects in spiral artery remodeling result in placental insufficiency and fetal distress. **A,** Schematic of workflow for Doppler ultrasound of pregnant dams. The embryo, UA/UV (yellow arrow), and placenta (dotted white outline) are labeled. **B,** Representative umbilical artery Doppler waveforms from two control and two Cyp19^Cre^; Cdh5^fl/fl^ placentas. The placenta is outlined in a dotted white line. Reversal of end-diastolic flow is evident by the change of directional velocity at the end of diastole compared to peak systole (i.e., negative to positive velocity). **C-E,** Quantification of resistance index (RI), pulsatility index (PI), and fetal heart rate. PSV (peak systolic velocity), EDV (end diastolic velocity), UA/UV (umbilical artery/umbilical vein). Note that red/blue colors in color Doppler images do not indicate UA/UV, which can only be differentiated based on the Doppler waveform. Control n=10, Cyp19^Cre^; Cdh5^fl/fl^ n=4. Statistical analysis was performed using two-tailed, unpaired Welch’s t-test. Data are shown as means±S.D.

## Discussion

The invasion of the maternal decidua by trophoblasts and their fusion to maternal SAs is a critical step in establishing placental circulation. However, the mechanisms by which trophoblast migration and endovascular invasion are accomplished remain largely unknown. Vascular mimicry describes a model in which trophoblasts express endothelial molecular and genetic programs^4^. However the exact function of specific endothelial genes in trophoblasts has not been functionally assessed in vivo, and this model remains untested. In blood and lymphatic ECs, VE-cadherin is used to maintain vascular integrity^13–15^, restrict endothelial migration^14,20^, and regulate angiogenic growth^21^. While it is an attractive concept that trophoblasts may form vascular sinuses using similar genetic programs, the findings that loss of VE-cadherin decreases trophoblast cell migration and prevents SA remodeling suggests that trophoblasts utilize VE-cadherin in a manner distinct from ECs. Our first genetic test of vascular mimicry in the placenta suggests that the use of endothelial proteins by trophoblasts may be relatively specific to their role in the placenta and not a simple reflection of vascular EC function.

In humans, defective SA remodeling and shallow trophoblast invasion are hallmarks of preeclampsia. Preeclampsia is a complex and heterogeneous disease with maternal and fetal contributions to its pathogenesis, and many in vitro models fail to fully recapitulate many aspects of its pathophysiology. Most rodent models of preeclampsia utilize maternal genetic or pharmacologic perturbations^22,23^, and there have been few in vivo models in which preeclamptic features are recapitulated with fetal modulation of trophoblasts. Previous studies of human placentas showed that invasive trophoblasts in severely preeclamptic placentas exhibit reduced expression of VE-cadherin^11^, however whether this is a cause or consequence of placental dysfunction has been unclear. Our mouse model utilizing trophoblast-specific knockout of VE-cadherin exhibits many histopathologic and clinical features of preeclampsia and suggests that loss of VE-cadherin in trophoblasts may be a primary contributor to preeclampsia pathogenesis. Furthermore, trophoblast-specific loss of VE-cadherin may serve as a useful model for studying fetal contributions to preeclampsia.

## Acknowledgments

We thank members of the Kahn laboratory for thoughtful discussions during the course of these studies, the CDB Microscopy Core for support with microscopy, and the Small Animal Imaging Facility Core for their support with ultrasound imaging. We thank Dr. Jeremy Veenstra-Vanderweele (Columbia University) and Dr. Gustsavo Leone (Medical University of South Carolina) for kindly providing the Cyp19^Cre^ mice.

## Author Contributions

DCS designed and performed most of the mouse and tissue culture experiments. XC, MC, YX, SG, and TCSK contributed to mouse studies. JY performed histological studies. SS performed Doppler ultrasounds. AB performed bioinformatics analysis. ML provided animal husbandry support. YY and JPS generated and characterized the Cdh5 conditional allele. DCS and MLK interpreted data and wrote the manuscript.

## Data and Materials Availability

All data and reagents will be made available upon reasonable request. Transgenic mouse lines not available through public repositories are available from Mark Kahn under a material transfer agreement with the University of Pennsylvania.

## Methods

### Generation of Mutant Mice

Cyp19(Tg)-Cre mice have been previously described^17^ in which the transgene relies on Cre expression under the Cyp19 promoter containing regulatory elements for trophoblast-specific expression, as Cyp19 is not endogenously expressed in trophoblasts. VE-cadherin (Cdh5) floxed mice have been previously described^24^ and were generated with LoxP sites flanking Exons 3 and 4. Mice were bred according to standard protocols and maintained on a mixed background. Male Cdh5^fl/fl^ mice were mated to female Cyp19^Cre^; Cdh5^fl/+^ mice due to the influence of parental inheritance on Cre expression, with maternal inheritance providing the most robust and consistent expression^17^. Mating pairs were set up in the afternoon and vaginal plugs checked in the morning. Presence of a vaginal plug indicated embryonic day (E)0.5. Cre-negative (Cdh5^fl/+^ and Cdh5^fl/fl^) and Cre-positive heterozygous (Cyp19^Cre^; Cdh5^fl/+^) littermates were used as controls. All procedures were conducted using an approved animal protocol (806811) in accordance with the University of Pennsylvania Institutional Animal Care and Use Committee.

### Intrauterine Doppler Ultrasound

In utero Doppler ultrasound was performed by a trained technician using the VEVO2100 Ultrasound System equipped with the MS-400 transducer (30MHz). E12.5 pregnant mice were lightly anesthetized using 2% isoflurane. Hair was removed from the abdomen using chemical hair remover (Nair), and the animals were placed on a warming pad. Maternal heart rate and temperature were continuously monitored and consistently within 400-500 bpm and 37°C. Ultrasound gel was applied to the abdomen and the transducer applied to visualize embryos and placentas using the maternal bladder as an anatomical landmark. Color Doppler was used to visualize the umbilical vessels, and pulse wave (angle of insonation <60°) measurements were made at the point where the umbilical artery inserts into the placenta. From the Doppler waveforms, peak systolic velocity (PSV) and end diastolic velocity (EDV) were measured and used to calculate resistance index [RI = 1 – PSV/EDV] and pulsatility index [PI = (PSV – EDV)/mean velocity, where mean velocity = (PSV+EDV)/2]. Heart rate was calculated as beats per minute by dividing 60 seconds by the systolic+diastolic time.

### Histology and Immunofluorescence Staining and Analysis

Whole mouse embryos or placentas were collected and fixed in 4% paraformaldehyde (PFA) overnight at 4°C prior to dehydration in alcohol and paraffin embedding. Tissue sections underwent to dewaxing and rehydration through xylene and ethanol treatment and were then subject to hematoxylin and eosin (H&E) staining or processed for immunofluorescence. For immunodetection, 10mM citrate buffer (pH 6) was used for antigen retrieval, and sections were blocked with 10% donkey serum in 1% BSA prior to primary antibody treatment overnight at 4°C. A list of antibodies can be found below. Fluorescence-conjugated Alexa Fluor secondary antibodies were used (1:500, Invitrogen) according to the primary antibody species and counterstained with DAPI (1:1000). Sections or tissues were mounted on slides with ProLong Gold Antifade reagent. Signals were detected and images collected using a Zeiss LSM 880 confocal microscope and Zeiss Axio Observer 7 widefield microscope. Images were visualized using ImageJ/FIJI software (NIH).

### Quantification of Immunofluorescence Images

Number of CK8^+^ trophoblasts and aSMA^+^ smooth muscle cells were manually counted and divided by total decidual area. Trophoblast invasion depth was measured as the distance the farthest CK8^+^ trophoblast was found in the decidua relative to the junctional zone. Percent labyrinth endothelial cell coverage was measured by quantifying Endomucin-positive area as a percentage of total labyrinth area. Trophoblast-endothelial displacement was quantified by measuring the circumference of the vessel lumen and then measuring the length of CK8^+^ trophoblasts in contact with the lumen. Percent displacement was calculated by dividing the trophoblast length by the lumen circumference. All images were analyzed using ImageJ/FIJI software.

### Antibodies

Antibodies for immunofluorescence: Endomucin (1:400, R&D AF4666 or 1:300, Abcam ab106100), TER119 (1:300, Abcam ab91113), mouse VE-cadherin (1:300, R&D AF1002), CK8 (1:300, Abcam ab53280 and 1:400, DSHB TROMA-1), αSMA-Cy3 (1:300, Sigma XXX)

### Statistical Analysis

All data are reported as means with n≥3 independent experiments or mice, and error bars represent standard deviation (SD). Each data point in the figures represents one individual placental or embryo. The explicit number of samples is indicated in the figure legends. No explicit power analyses were used to predetermine sample size, and no randomization was used. No samples were excluded for analysis. Statistical significance was determined using Welch’s t-test. Differences between means were considered significant at P<0.05. Significant differences in expected genotypes was calculated using Fisher’s exact test and considered statistically significant at P<0.05.

**Figure 1-Table Supplement 1.** Decreased survival of Cyp19^Cre^; Cdh5^fl/fl^ mutants in late gestation. Cdh5^fl/fl^ male mice were crossed with Cyp19^Cre^; Cdh5^fl/+^ females to generate litters with mixed genotypes. The expected percentage is listed under the genotype label. The observed number of each genotype is shown with the corresponding percentage given in parentheses. P-values were calculated using Fisher’s exact test at stages E10.5-12.5 (pooled), E14.5-16.5 (pooled), and P21. E designates embryonic day and P designates postnatal day. *One embryo at E12.5 was found to be dead.

|  | **Cyp19<sup>Cre</sup>; Cdh5<sup>fl/fl</sup>**<br>(25%) | **Cyp19<sup>Cre</sup>; Cdh5<sup>fl/+</sup>**<br>(25%) | **Cdh5<sup>fl/fl</sup>**<br>(25%) | **Cdh5<sup>fl/+</sup>**<br>(25%) | **Total**<br>(100%) | **Fisher Exact**<br>**Test** |
| --- | --- | --- | --- | --- | --- | --- |
| **E10.5-E12.5** | 12* (26.7%) | 12 (26.7%) | 9 (20%) | 12 (26.7%) | 45 (100%) | *P*=0.7667 |
| **E14.5-E16.5** | 1 (2.9%) | 13 (38.2%) | 12 (35.3%) | 8 (23.5%) | 34 (100%) | *P*=0.0033 |
| **P21** | 1 (2.2%) | 11 (24.4%) | 15 (33.3%) | 18 (40.0%) | 45 (100%) | *P*=0.0333 |

**Figure 1-Figure Supplement 1.**
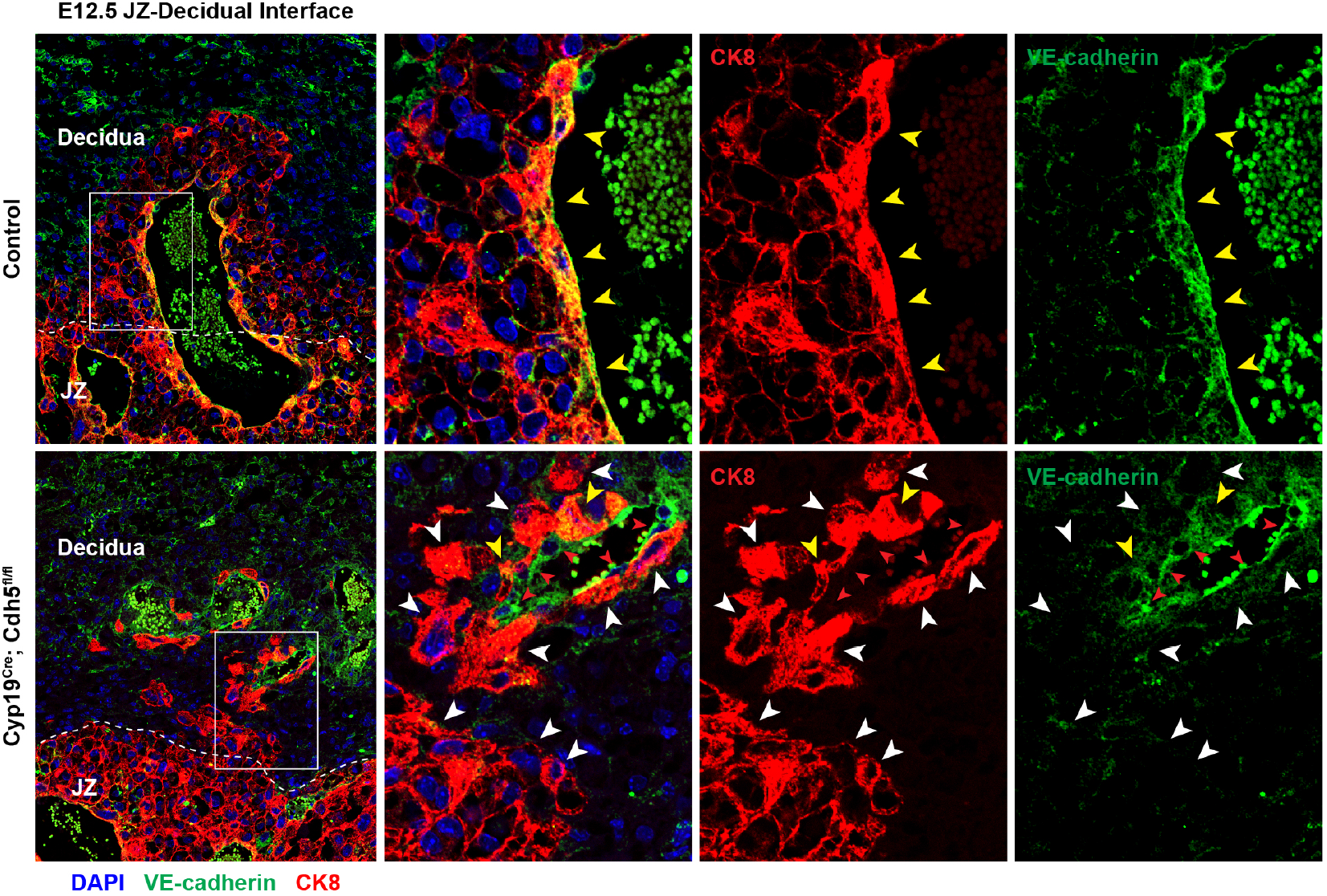
Deletion of VE-cadherin in Cyp19^Cre^; Cdh5^fl/fl^ placentas. Immunofluorescence staining of E12.5 Control and Cyp19^Cre^; Cdh5^fl/fl^ placentas for VE-cadherin (green) and CK8 (red). Yellow arrowheads indicate VE-cadherin^+^ trophoblasts, which predominantly line remodeled spiral arteries. White arrowheads indicate VE-cadherin^-^ trophoblasts. Red arrowheads indicate spiral artery ECs, which are VE-cadherin^+^ CK8^-^. Note that there are a few VE-cadherin^+^ trophoblasts in Cyp19^Cre^; Cdh5^fl/fl^ placentas, however VE-cadherin is deleted from the majority of trophoblasts. The layer of VE-cadherin^+^CK8^-^ cells (red arrowheads) in the Cyp19^Cre^; Cdh5^fl/fl^ placentas are spiral artery endothelial cells that have not been displaced by trophoblasts. Dotted white lines demarcate the decidua and junctional zone (JZ).

**Figure 1-Figure Supplement 2.**
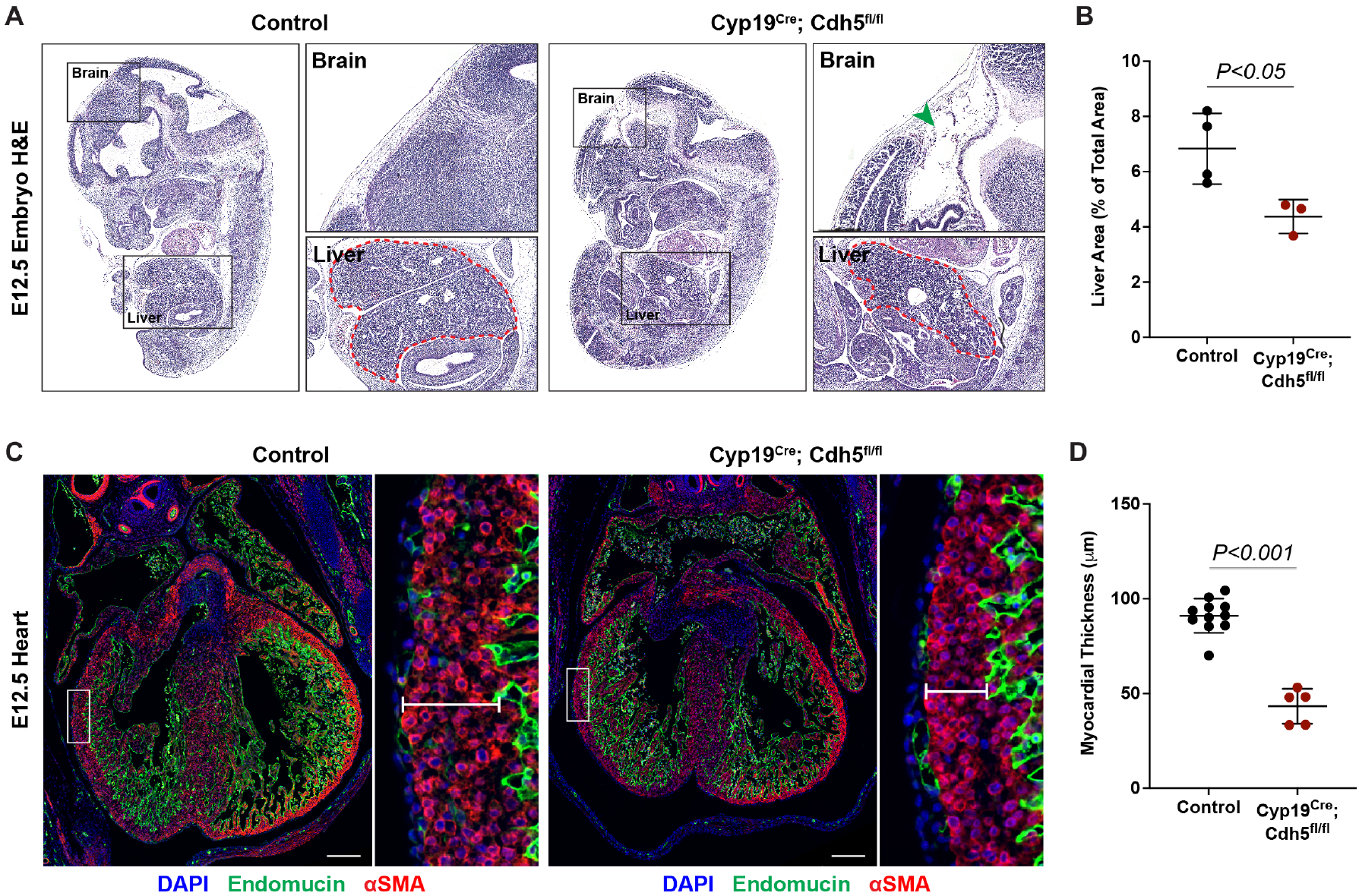
Loss of trophoblast VE-cadherin causes defects in brain, liver, and heart development. **A,** Sagittal H&E sections from E12.5 control and Cyp19^Cre^; Cdh5^fl/fl^ embryos. Boxes show magnified regions of the brain, heart, and liver. Note the thinness of the myocardium in Cyp19^Cre^; Cdh5^fl/fl^ embryos. Dotted red line outlines the liver contour. Green arrowhead points to tissue degradation in the brains of Cyp19^Cre^; Cdh5^fl/fl^ embryos. **B,** Quantification of the liver area as a percent of total embryo area. Control n=4, Cyp19^Cre^; Cdh5^fl/fl^ n=3. **C, D,** Immunofluorescence staining of E12.5 control and Cyp19^Cre^; Cdh5^fl/fl^ embryonic hearts for Endomucin (green) and aSMA (red). Insets demonstrate myocardial thinning and quantified in (**D**). Control n=11, Cyp19^Cre^; Cdh5^fl/fl^ n=5. Scale bars = 200μm. Statistical analysis was performed using two-tailed, unpaired Welch’s t-test. Data are shown as means±S.D.

